# Inferring Cell–Cell Interaction Dynamics from Cell Trajectory Data Using Deep Attention Networks

**DOI:** 10.64898/2026.07.14.707033

**Authors:** James Boyle, Helen M Byrne, Ruth E Baker

## Abstract

Interactions between nearby cells are a key driver of cell movement in many biological systems, including collective cell migration and the immune response to cancer. However, inferring the interaction rules in a given system in a manner that is both accurate and biologically interpretable remains a challenge. A valuable experimental method for analysing cell–cell interaction dynamics is the tracking of individual cell locations over a series of time-lapse images, and in this work we present a model, based on the theory of deep attention networks, that learns how cell–cell interactions affect cell movement directly from cell trajectory data. Our approach requires no *a priori* assumptions about the mechanisms governing cell behaviour, enabling it’s application to cell trajectory data originating from a diverse range of biological systems.

In addition to the model, we develop a suite of tools that exploit the model’s attention-based structure to present the learned interaction dynamics in an interpretable manner. Our model extends previous applications of deep attention networks to cell movement by providing deeper insights into cell–cell interaction dynamics, moving beyond inferring whether cells interact to inferring how these interactions affect cell movement, and providing the ability to infer type dependent cell–cell interaction dynamics in multi-type cell movement systems. By combining data-driven learning and structural interpretability, our approach represents a highly general methodology for linking cell trajectory data to mechanistic hypotheses, showing that deep attention networks constitute a powerful exploratory tool for characterising the effect of cell–cell interactions on cell movement in complex cellular systems.

## 1 Introduction

Cell movement is an important component of many biological and biomedical processes, including embryonic development [1], collective cell migration [2, 3], tumour invasion and metastasis [4], and the immune response to cancer [5, 6]. In many such systems, a key driver of cell movement is interactions between spatially proximal cells. Interactions may be short range, such as intracellular responses to cell contact [7–10], or forces arising from cell–cell adhesion [7, 11, 12], friction [7, 13, 14] or repulsion [7, 11]. Cells also interact over longer distances [15]; this can include direct contact involving elongated cell protrusions such a pseudopodia [16, 17], or interactions mediated by a secondary agent, such as chemical signalling [9, 15, 18], or the transmission of physical forces through the extracellular matrix [15]. In order to fully understand how cells move in a given system, we thus must understand how local and non-local cell–cell interactions affect cell movement.

Advances in live cell imaging technologies and cell tracking algorithms have made it possible to collect individual cell trajectory data [19, 20]. This consists of the co-ordinates of each cell in an assay over a series of time points, typically computed by applying a cell tracking algorithm to a series of time-lapse images in which individual cells are distinguishable [21] (figure 1A). Compared to bulk measurements of cell migration, single-cell trajectories provide more detailed information on the factors influencing cell movement, and on how cells integrate information from their local environment into movement behaviours. Realising the full potential of these datasets requires methods that can directly infer the effects of cell–cell interactions on cell movement from observed cell trajectories.

**Figure 1:**
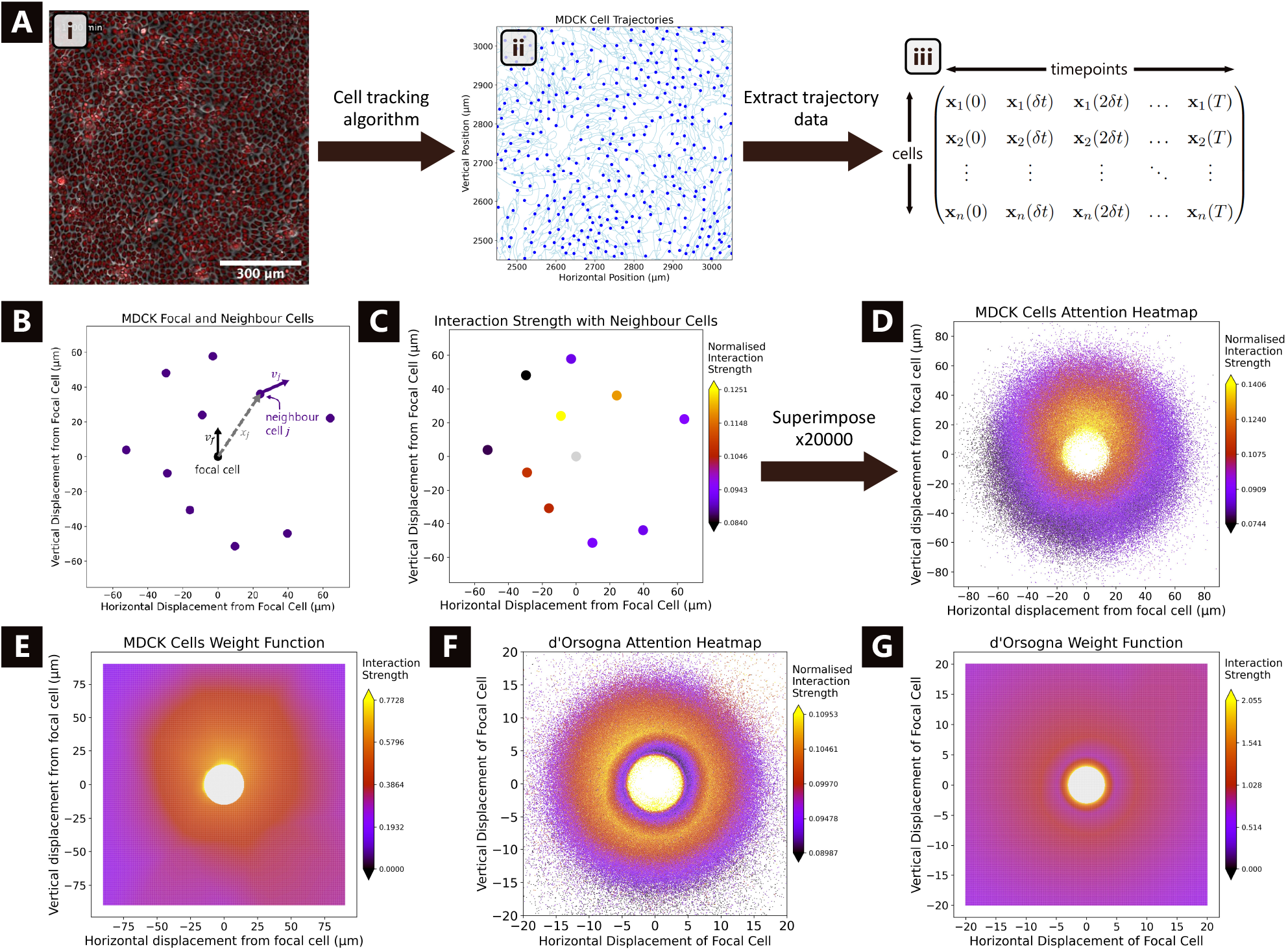
Generating an attention heatmap from trajectory data. A) Outline of the process of obtaining cell trajectory data: i) Snapshot of collectively migrating MDCK; ii) Cell tracks for a subset of the MDCK cells. Blue dots are the end locations of each cell, with trajectories in light grey lines behind them; iii) Matrix containing the location of each cell at each time point in the data. This is the data a deep attention network sees during training. B) A single focal cell (in black) and its 10 nearest neighbour cells at a single time point. C) Focal and neighbour cells from panel B, with neighbour cells coloured by their influence on the focal cell’s movement, as determined by the weight function from a trained deep attention network. D) Attention heatmap for the MDCK cells, produced by the superposition of plots like panel C for 20,000 different focal cells at different time points. E) Plot of the trained weight function for the MDCK cells. F) Attention heatmap for the simulated data from the d’Orsogna model. G) Plot of the trained weight function for the d’Orsogna data. Figure A(i) is reproduced from [28] under the terms of the Creative Commons Attribution License.

In this paper we present a new deep-learning model, based on the theory of deep attention networks, that infers how cell–cell interactions affect cell movement directly from cell trajectory data [22–24]. The expressivity and function learning capabilities of deep neural networks [25–27] enable cell–cell interaction dynamics to be learned in a hypothesis-free manner, and the model architecture enables the learned dynamics to be automatically presented in a biologically interpretable format. This ability to unbiasedly learn and then automatically present cell–cell interaction dynamics makes the model a powerful exploratory tool for studying cell movement.

Deep attention networks have previously been used to analyse cell trajectory data [28, 29], as well as the migration of schools of zebrafish [30, 31]. Existing deep attention network models have focused on predicting the binary turning behaviour of cells (whether a cell will turn left or right during a given time period), so can only learn cell–cell interactions that significantly affect this discrete output. The deep attention network model developed in this paper is trained to predict a cell’s complete movement vector; in order to do this, it must learn the full range of cell–cell interaction dynamics that affect cell movement. This change in predictive target enables the trained deep attention network to infer not only how strongly cells affect their neighbours’ movement, as in previous applications of deep attention networks to cell movement data [28, 29], but also how these interactions alter cell movement, providing deeper insight into the mechanisms of cell motility. A further significant conceptual development is the extension of the deep attention network architecture to learn type-dependent cell–cell interaction dynamics in systems comprising multiple motile cell types.

The remainder of the paper is structured as follows. In section 2 we motivate and define the new deep attention network model, before showing how it can be used to determine the relationship between cell displacement and interaction strength in an assay of collectively migrating Madin–Darby Canine Kidney (MDCK) epithelial cells. These well-studied cells move as a highly coordinated, tightly packed collective, producing characteristic swirling domains [28]. A natural question is how individual MDCK cells respond to information from their local neighbourhood to make the movement decisions that give rise to this collective behaviour. Our deep attention network analysis suggests that MDCK cells take most of their movement cues from their frontward neighbours. The MDCK cell movement assay we use was first analysed using deep attention networks in [28], and in section 3, we present a direct comparison between that implementation and our model, demonstrating the effect of the changes we introduce. In section 4 we show how our deep attention network model provides richer insight into the mechanisms of cell motility by inferring the downstream effect of cell–cell interactions on cell movement. In particular we use the deep attention network model to infer the presence of cell–cell attraction and repulsion, and velocity alignment, in the same MDCK cell assay. Finally, in section 5 we show how the deep attention network architecture may be extended to infer cell-type dependent interaction dynamics in systems with multiple motile cell types.

## 2 Deep attention networks

In order to infer cell–cell interaction dynamics directly from cell trajectory data, we assume we know the location of *n* cells at regularly spaced time points *t* = *δt*, 2*δt*, …, *Kδt* (figure 1A(iii)). Ideally, these data should include all cells present in the experiment (i.e. no cells should be untracked), and each cell should come close enough to interact with other cells for at least some portion of the experiment. We aim to use these data to understand how a cell’s movement is affected by its interactions with spatially proximal cells. To do this, we train a model to predict where a cell will move over a specified time interval based on its velocity, and the displacement and velocity of nearby cells, at the beginning of the interval. More precisely, if we take a single cell (the “focal cell”) at some time *t*, we aim to predict the cell’s movement vector over a fixed time interval [*t, t* + Δ*t*), denoted by Δ**x** (if **X**(*t*) denotes the focal cell’s position at time *t*, then Δ**x** = **X**(*t* + Δ*t*) − **X**(*t*)). We aim to predict Δ**x** as a function of the focal cell’s velocity **v**_*f*_ at time *t*, as well as the velocity (**v**_*j*_) and relative displacement (**x**_*j*_) of its *N* nearest neighbour cells *j* = 1, …, *N* at time *t* (figure 1B), where both *N* and Δ*t* are fixed in advance.

We decompose the prediction of Δ**x** into a sum of terms, each arising from the interaction between the focal cell and a single neighbour cell, i.e. 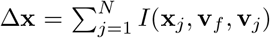 for some interaction function *I*. Each of these terms is then decomposed further into two parts. The first is a vector valued function Π(**x**_*j*_, **v**_*f*_, **v**_*j*_) that measures the contribution to the focal cell’s movement resulting from its interaction with neighbour cell *j*, based on the relative displacement of neighbour cell *j* and the velocities of the focal cell and neighbour cell *j*. The second is a positive real-valued function *W* (**x**_*j*_) that measures the the relative influence that neighbour cell *j* has on the motion of the focal cell, based on the displacement of the neighbour cell. The focal cell’s movement is thus predicted as

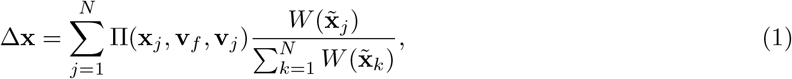

where the tilde indicates that the input to *W* is given in co-ordinates where the focal cell’s velocity is aligned with the positive *y*-axis.

The function Π is called the *pairwise interaction function* and *W* is called the *weight function*. The output of *W* is most properly interpreted as a statistical measure of the relative influence that a given neighbour cell has over the movement of the focal cell, but may also be more succinctly referred to as measuring the “interaction strength” between the focal and neighbour cell [28, 29]. The final prediction of Δ**x** is a weighted average of the pairwise interaction terms, with the weights determined by the weight function.

Both Π and *W* are fully connected deep neural networks, allowing the exact cell–cell interaction dynamics to be learned directly from data in a hypothesis-free manner. Neural networks are trained using the standard combination of backpropogation and adaptive moment estimation—see the supplementary material for details on network architecture and training. For the remainder of this paper, we shall refer to the model defined by equation (1) as a *Deep Attention Network*.

The strength of the deep attention network model lies in its *interpretability*. In particular, the additive nature of equation (1), and the decomposition of each interaction term into separate functions for measuring the effect and strength of an interaction, allow for the application of various model interpretability techniques to infer how cell–cell interaction dynamics depend on the relative displacement of the interacting cells.

### 2.1 Attention heatmaps

Once we have trained a deep attention network on a cell trajectory dataset, we can use the trained weight function to determine how strongly cell movement is affected by different neighbouring cells. To illustrate this, we trained a deep attention network on cell trajectory data from an assay of collectively migrating MDCK cells, a line of kidney epithelial cells [28]. The MDCK cells move in a highly coordinated fashion, producing characteristic swirling domains [28] (see figure 1A(i) for a snapshot, and [28] for the full video and more detailed information).

Figure 1C shows a single MDCK cell at the centre, and its *N* = 10 nearest neighbour cells at a given time point, with each neighbour cell coloured by the normalised strength of its interaction with the focal cell (i.e. the colour of neighbour cell *j* is determined by 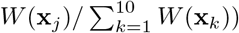. In this instance the strongest interaction is with the neighbour cell almost directly in front of the focal cell. We gain a fuller understanding of the effect of cell–cell displacement on interaction strength by superimposing thousands of such plots for multiple focal cells at different time points. For example, figure 1D shows that, in this data set, the movement of MDCK cells is most strongly influenced by neighbouring cells to the front and immediately behind, with interaction strength decaying as one moves further away and further behind the focal cell.

We call the plot in figure 1D an *attention heatmap* and, following [28, 29], we say that *W* measures how much “attention” a focal cell pays to its neighbour cells [28]. We emphasise that the term “attention” comes from the deep learning literature^1^, and does not necessarily reflect any “intentional” or specific form of cell–cell interaction. For example, if a cell’s movement is physically impeded by a cell in front, then the impeding cell exerts a strong influence on the impeded cell’s movement; in this case we would say that the impeded cell “pays attention” to the impeding cell.

As the learned weight function depends only on the displacement of the neighbour cell, we can plot it directly. The learned weight function for the MDCK cells is plotted in figure 1E, and shows the same forward bias in attention present in the attention heatmap. The unnormalised weight function can be a useful complementary inference tool, but should be used with care. The deep attention network is only trained to produce accurate predictions when applied simultaneously to multiple neighbour cells, with the distribution of displacements and velocities found in the training data. For example, in figure 1E we have not plotted the weight function within a small radius of the origin because, as can be seen from the attention heatmap, there were no instances in the trajectory data where cells were within that distance of each other. Using the attention heatmap ensures that the trained deep attention network is only evaluated in the region of input space on which it has been trained.

### 2.2 Simulated data

To verify that patterns in attention heatmaps are reflective of genuine dynamics in the underlying system, we trained deep attention networks on simulated cell trajectory data, generated using agent-based models of cell movement. Using agent-based models we can specify exactly how the movement of each cell is affected by its neighbours, providing a “ground truth” to which we can compare results obtained from a trained deep attention network.

Here we present the results of training a deep attention network on simulated data from the well-studied *d’Orsogna* model [32], in which cell movement is affected by persistence, noise, and proximity-dependent forces exerted by neighbouring cells. The forces are defined such that cells strongly repel each other at short distances, and weakly attract each other at medium distances, with an intermediate distance at which cells neither attract nor repel each other (see the supplementary material for model equations). For the *in silico* experiments analysed in this section, we simulated 2000 cells initially seeded at random in a circle of radius 150 (in this paper, all quantities relating to simulated data are non-dimensional), with the repulsive force fully dissipating at a distance of approximately 4 and an attractive zone centred at a distance of approximately 4.8 between interacting cells (model parameters are given in the supplementary material). Figure 1F,G shows the attention heatmap and unnormalised weight function from the trained deep attention network. Zones of strong and weak interaction at short and medium distances are clearly visible. More details on the results of training deep attention networks on this data, can be found in the supplementary material.

## 3 Relationship to previous work

As discussed in the introduction, structurally similar models to (1) have been used to analyse cell trajectory data [28, 29]. Indeed the notion of an attention heatmap^2^ was first introduced in [28]. The model presented in equation (1) represents the result of a series of developments implemented in order to increase the interpretability and reliability of inferences made using the trained deep attention network. The aim of these changes was to make deep attention networks a more reliable general-purpose tool for exploratory analyses of cell trajectory data, producing results that more accurately reflect the underlying biology, and are less sensitive to artefacts surrounding experimental setup or model specification. In this section we describe these changes, and provide a detailed comparison of results obtained using equation (1) and the deep attention network model used in [28–31]. This section can be skipped by readers wishing only to understand the inferential capabilities of the model in equation (1).

The model used in [28, 29] is structurally similar to equation (1), but instead aims to predict the log-odds^3^ *z* that the focal cell will turn right (as opposed to turning left) over [*t, t* + Δ*t*) as

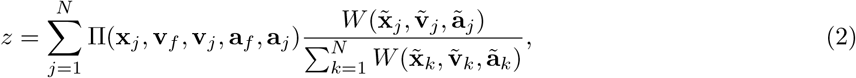

where **a**_*f*_, **a**_*j*_ are the accelerations of the focal cell and neighbour cell *j* (all other variables are as before, and tildes continue to indicate variables expressed in coordinates with the *y*-axis aligned with the focal cell’s velocity). The functions Π and *W* are again fully connected neural networks with the same hidden layers.

The most significant difference between equation (2) and the model we propose in equation (1), is that we aim to predict the focal cell’s full movement vector over [*t, t* + Δ*t*), rather than just the binary direction in which it will turn. This enables the trained model to learn a wider range of cell–cell interaction dynamics, as many interactions can significantly affect the focal cell’s movement without having a dramatic impact on its turning behaviour. Moreover, displacement is a continuous and more robust prediction target. Small changes in the focal cell’s movement will result in small changes to its displacement, but could alter whether it turns left or right (conversely, very large changes in focal cell movement can result in no change to its turning behaviour; see supplementary figure 1).

Secondly, we constrain the weight function to depend on position only. This is motivated by the fact that *W* is predominantly used to infer the dependence of cell–cell interaction strength on relative displacement. It also allows the weight function to be plotted directly. We also remove the dependence on acceleration of the weight and pairwise interaction functions, motivated by experiments in [28] showing its inclusion does not significantly affect the learned dynamics.

Finally, in equation (2) symmetry requirements are imposed on Π and *W* . Π is constrained to be anti-symmetric with respect to reflection in the *y*-axis, and inputs to *W* are given in a *y*-reflection invariant form, so that *W* is symmetric with respect to reflection in the *y*-axis [30]. These constraints restrict the possible parameter space for Π and *W*, and were motivated by the need to enable to overall model to be trained on the relatively low amount of data typically available from observations of animal movement [30]. As cell movement assays typically produce much larger amounts of data, a deep learning model should be able to learn these symmetries if they are present. Moreover, the introduction of hard post-hoc constraints such as these can hinder the training of neural networks [33–35]. Further, when considering cell movement, we would not expect the interaction dynamics to necessarily have these symmetries. Therefore, in (1) we remove these symmetry requirements and allow Π and *W* to be fully unconstrained neural networks.

For ease of reference, for the rest of this section we refer to the model in equation (1) as a *generalised* deep attention network (gDAN), and the model in equation (2) as a *turn* deep attention network (tDAN). To evaluate the impact of the differences between them, we compare inferences from a tDAN and gDAN trained on trajectory data from the MDCK cell assay discussed above. Recall that this assay was first analysed using a tDAN in [28]. In [28], cells were split into “edge” cells (those further than 80% of the tissue radius from the tissue centre) and “bulk” cells (all remaining cells)^4^.

Figure 2A shows attention heatmaps for the MDCK edge cells produced by successively making each of the changes described above (removal of symmetry requirements, predicting movement instead of turning behaviour, and removal of velocity dependence from the weight function) to the tDAN model. Both models trained to predict turning behaviour produce attention heatmaps with distinctive “fans” of attention to the front left and front right of the focal cell, and a void in attention to the rear of the focal cell. These features are widely observed in tDAN attention heatmaps [28–31], and have motivated claims of anisotropy and directional polarity in cell neighbour sensing [29]. We hypothesise that this form of heterogeneity in attention does not come from the underlying biology, but rather from which cell–cell interactions most strongly affect the binary “turn left/turn right” flag. This is physically intuitive: a neighbour cell to the front right (or front left) of the focal cell is likely to strongly influence whether the focal cell turns right or left by physically impeding it from turning in one direction. That these features do not appear in the attention heatmaps from models that predict general displacement supports the hypothesis that these features are an artefact of the choice of predictive target. Figure 2A also demonstrates that removing velocity dependence from the weight function produces sharper attention heatmaps that are easier to interpret.

**Figure 2:**
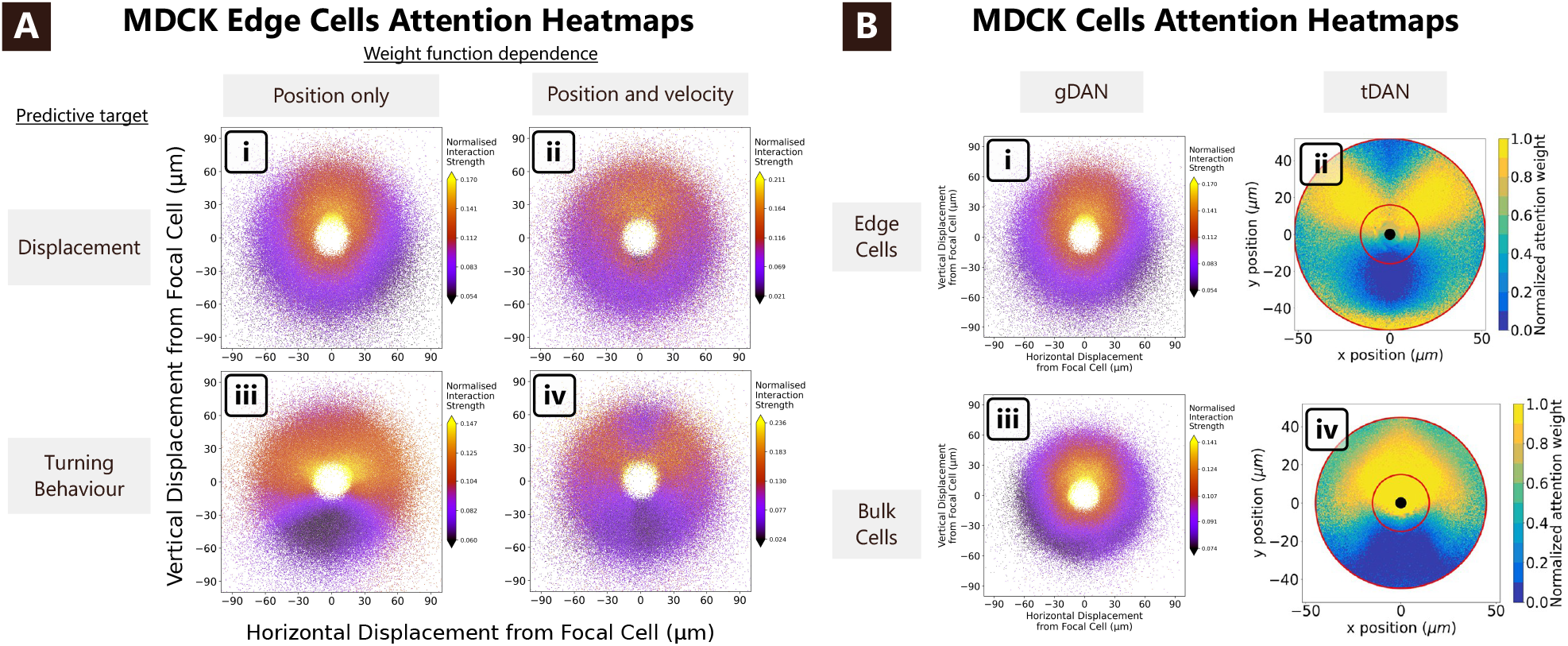
A) Attention heatmaps produced by different deep attention network models trained on the edge MDCK cell trajectory data. Rows indicate what the model was trained to predict, and columns indicate which variables were included in the weight function (so for (i) and (iii) *W* = *W* (**x**_*j*_), and for (ii) and (iv) *W* = *W* (**x**_*j*_, **v**_*f*_, **v**_*j*_)). Panel (iv) corresponds to a tDAN with the symmetry constraints removed, whilst panel (i) corresponds to a gDAN. B) Attention heatmaps from gDAN and tDAN models trained on bulk and edge MDCK cells. The gDAN produces more consistent results for the edge and bulk cells. Figures B(ii) and B(iv) are reproduced from [28] under the terms of the Creative Commons Attribution License

Figure 2B shows attention heatmaps produced by training a tDAN and a gDAN on trajectory data from the populations of bulk and edge cells. These are the same cell line, so to the extent that the attention heatmaps reflect underlying biology, we would expect them not to vary much between the two populations. The tDAN attention heatmaps vary substantially between the two populations, whereas the gDAN attention heatmaps are more consistent, yielding qualitatively similar mechanistic conclusions. The gDAN attention heatmaps thus appear more robust to differences in experimental conditions—here the differences between the bulk and edge of the expanding monolayer. Nevertheless the changes in biophysical conditions do induce some variation in the gDAN attention heatmaps, so care should be taken in considering how local factors such as cell density may affect patterns of attention. In this case, edge cells occupy a less cell-dense environment, particularly ahead of them, making it more likely that “further ahead” neighbour cells exert a stronger influence on future movement, since fewer nearby cells are present to dominate the focal cell’s dynamics.

Inference from a tDAN often requires substantial knowledge of how the specific details of the predictive model affect the results obtained from the trained deep attention network, hindering their use as an exploratory tool for analyses of cell movement data. If attention heatmaps are to be used to analyse cell trajectory data, practitioners should be confident that patterns in attention reflect the underlying biology, and not the result of experimental conditions or choices in the training of the underlying neural networks. To this end, the generalised deep attention network model in equation (1) represents significant progress towards establishing deep attention networks as a reliable exploratory tool for analysing cell trajectory data. For the remainder of the paper, whenever we talk about deep attention networks we shall refer to the model defined in equation (1).

## 4 Qualitative inference

Previous applications of deep attention networks to cell trajectory data have been limited to inferring which cells mostly strongly influence their neighbours’ movement. Beyond this, we would like to know *how* the interaction between two cells affects their movement. In this section we show how deep attention networks can be used to infer relationships between the relative displacement of interacting cells, and the effect of cell–cell interactions on cell movement.

This advance is enabled by the decomposition in equation (1) of the focal cell’s movement into a sum of terms, each representing the interaction with one neighbour cell. By removing a single neighbour cell from this sum (or equivalently, setting its weight to zero), we can compare where the focal cell will move both when a particular neighbour is present or absent, according to the dynamics learnt by the deep attention network. By comparing where the focal cell will move with and without the presence of a specific neighbour cell, we can infer the effect that the interaction with a given neighbour cell has on the focal cell’s movement.

Consider a single focal cell at some time *t*, with *N* nearest neighbour cells *j* = 1, …, *N* . Let

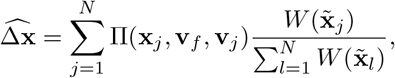

be the predicted location of the focal cell at time *t* + Δ*t* under the dynamics learnt by the deep attention network (recall we take the focal cell’s original location to be at the origin).

For a specific neighbour cell *k*, we estimate where the focal cell would move if neighbour cell *k* were not present by considering

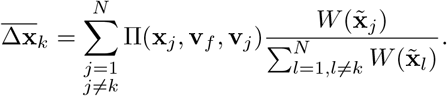

We then compare 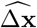 and 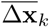 to determine how the focal cell’s movement is affected by its interaction with neighbour cell *k*.

### 4.1 Attraction–repulsion heatmaps

One thing we might wish to investigate is whether cells tend to attract or repel each other. We can inspect whether the interaction with neighbour cell *k* brings the focal cell nearer to, or further away from, the location of neighbour cell *k*. In this way, we can assign an “attraction score” to the interaction as

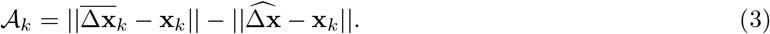

This measure will be positive if the interaction with neighbour cell *k* moves the focal cell moves closer to the location of the neighbour cell, *compared to the scenario when neighbour cell k is not present*, or equivalently when it has no effect on the focal cell’s movement (figure 3A).

**Figure 3:**
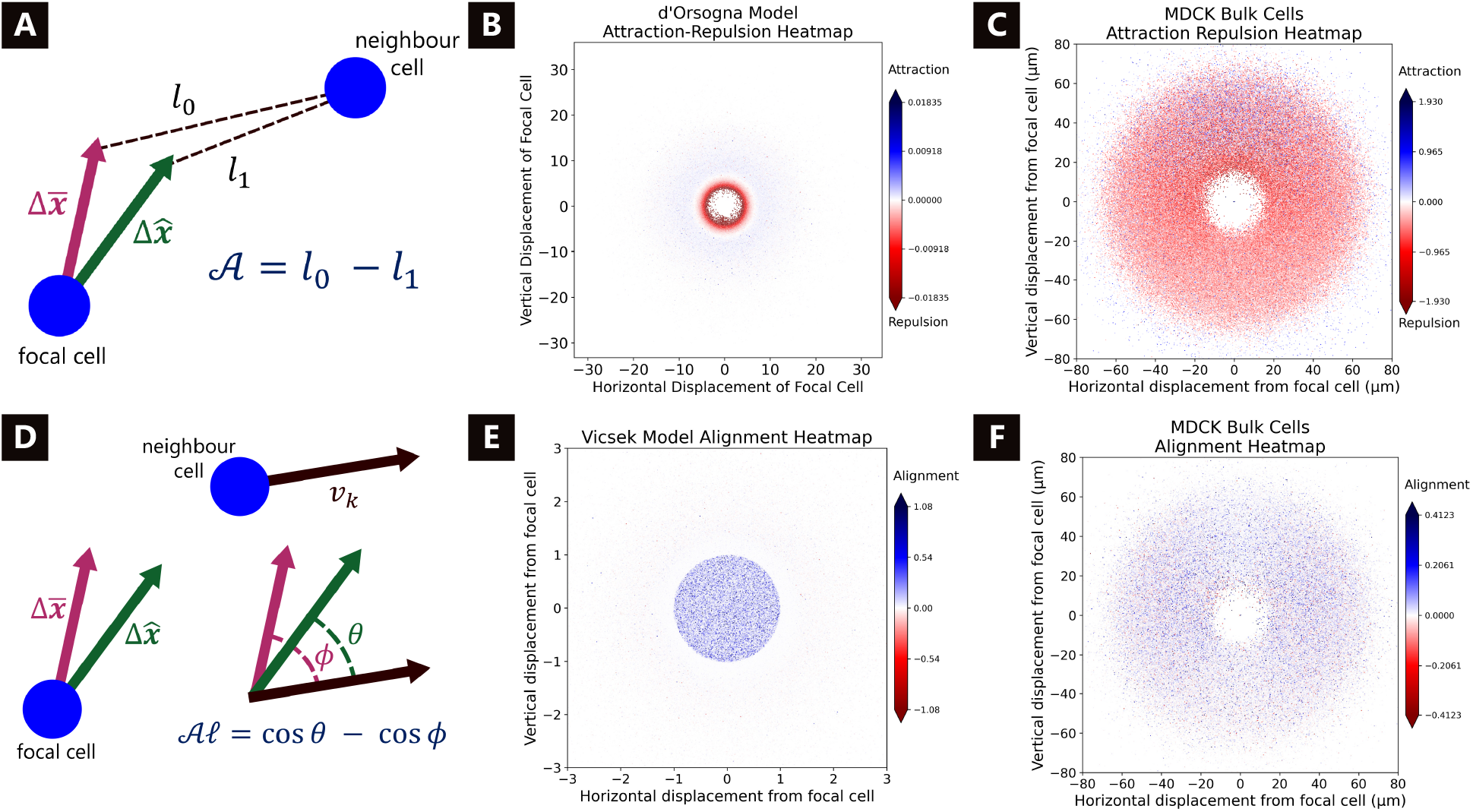
A) Illustration of the attraction score defined in equation (3): 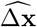 and 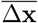 are the focal cell’s predicted movement vectors over Δ*t* with and without the neighbour cell present. B) Attraction–repulsion heatmap for the d’Orsogna model. C) Attraction–repulsion heatmap for the MDCK bulk cells. D) Illustration of the alignment score defined in equation (4). E) Alignment heatmap for the Vicsek model. F) Alignment heatmap for the MDCK bulk cells.

The attention heatmaps in figure 1 were produced by superimposing a large number of plots of neighbor cells coloured by 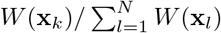, showing the dependence of cell–cell interaction strength on relative displacement. We can determine how displacement affects whether cells tend to attract or repel each other by instead colouring each neighbour cell by *A*_*k*_. Following previous nomenclature, we call such a plot an *attraction–repulsion heatmap*.

Figure 3B shows the attraction–repulsion heatmap for the simulated data from the d’Orsogna model discussed in section 2.2, in which cells interact by strongly repelling each other at short distances, and weakly attracting each other at medium distances. The attraction–repulsion heatmap recovers the zones of strong repulsion and weak attraction. The attention heatmap for this data (figure 1F) identified these zones as those with the strongest cell–cell interactions; the attraction–repulsion heatmap augments this with additional information about the nature of these interactions.

Figure 3C shows the attraction–repulsion heatmap for the MDCK bulk cells analysed previously. The heatmap shows that MDCK cells tend to be repelled by neighbouring cells to their side, rear, or immediately in front. Further in front (beyond approximately 30µm) a simple attractive or repulsive description based on location is insufficient to describe the full complexity of the cell–cell interactions in this region.

#### Interaction potential inference

When constructing mathematical models of cell movement, it is common to suppose cells interact according to an *interaction potential*. This is function *U* : [0, ∞) → ℝ, constructed such that the force between two cells separated by a distance *r* is given by *U* ^*′*^(*r*), with positive forces being attractive and *vice versa*. Accordingly, there is substantial interest in inferring interaction potentials from observations of cell movement. Fitting an interaction potential to data involves exactly determining how strongly cells attract and repel each other based on their relative displacement, and motivated by this we explored whether we could infer interaction potentials via radially averaging the trained weight function from a deep attention network.

In supplementary figure 2A we plot the potential *U* (*r*) and its derivative *U* ^*′*^(*r*) used to create the d’Orsogna simulated data analysed above. In supplementary figure 2B we plot the radially averaged weight function 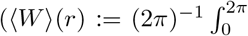 (*r* cos *θ, r* sin *θ*) d*θ*) from a deep attention network trained on this data. We augment this in supplementary figure 2C by plotting the normalised interaction strength for a large number of neighbour cells, coloured by their attraction score. We see that these plots capture many of the key qualitative features of the interaction potential, accurately identifying regions of strong repulsion and weak interaction at short and medium distances.

Whilst it is not immediately clear how one could extract a functional form for the interaction potential from a trained deep attention network, we note that in this example, radially averaging the trained weight function reveals many of the key qualitative features of how cell–cell interaction forces vary with distance. We also note that this is achieved without a pre-imposed hypothesis that an interaction potential will accurately model cell–cell interactions, and no information about the other mechanisms affecting cell movement. In this way, the trained weight function can also indicate whether an interaction potential is an appropriate model. For example, a non-radially symmetric weight function, as in figure 1E, would indicate that an interaction potential is insufficient to fully describe the interaction dynamics.

### 4.2 Alignment heatmaps

Another phenomenon commonly observed in cell migration assays is velocity alignment, in which nearby cells tend to align their velocities [7]. With the same setup and notation as above, we assign each neighbour cell *k* an “alignment score” as

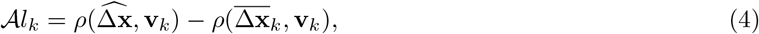

where *ρ*(**a, b**) = **a** · **b***/*∥**a**∥∥**b**∥ is cosine correlation (figure 3D). This score compares the orientation of the focal cell with and without the presence of neighbour cell *k*, and will be positive if the interaction with neighbour cell *k* causes the focal cell’s velocity to be more aligned with that of neighbour cell *k*, compared to if neighbour cell *k* were not present. As with attention heatmaps and attraction-repulsion heatmaps, we superpose plots of neighbour cells coloured by their alignment score to produce an *alignment heatmap*.

To validate that the alignment score and alignment heatmaps reflect genuine velocity alignment in the underlying cell movement system, we trained a deep attention network on simulated data from a Vicsek model [36]. In this model cells orient themselves to match the average orientation of their neighbours within a given perceptual zone (see the supplementary material for model details). We trained a deep attention network on data from this model in which the perceptual zone was a circle of unit radius around the focal cell (recall all quantities relating to simulated data are dimensionless, further details can be found in the supplementary material). This domain is clearly identified in the alignment heatmap (figure 3E). The alignment heatmap for the MDCK bulk cells (figure 3F) shows that they tend to align their velocities with that of neighbour cells in all directions, with this alignment being stronger for neighbour cells in front or to the side.

## 5 Cell movement in heterogenous cell populations

The model in equation (1) assumes homogeneous interaction dynamics across all cells. Yet many cell movement systems contain multiple motile cell types, which we would expect to influence each other’s movement in different ways. For example, migrating neural crest cells exhibit leader and follower phenotypes, with leader cells responding to external chemical gradients, and follower cells taking movement cues from leader cells [37, 38]. Additionally, in the immune response to invasion by a pathogen, different types of immune cells interact differently with each other and with pathogen cells [39].

In this section we extend our deep attention network model to analyse cell trajectory datasets with multiple cell types, in which the type of each cell is known *a priori*. We extend the model by including the types of the focal and neighbour cells as variables in the pairwise interaction and weight functions, so that the focal cell’s movement over [*t, t* + Δ*t*) is now predicted as

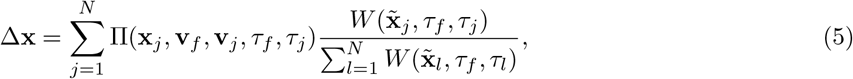

where *τ*_*f*_, *τ*_*j*_ are the types of the focal cell and neighbour cell *j*. We also increase the width and depth of the networks Π and *W* to allow them to learn more complex type-dependent interaction patterns (see the supplementary material).

The inference techniques described in previous sections extend naturally to the multiple cell-type setting.

For example, we can construct an attention heatmap showing how cells of type *A* pay attention to cells of type *B* by following the same procedure described in section 2.1, but only plotting cells of type *B* that are neighbours to a type *A* cell.

*Results from training a type aware deep attention network on a multi-type cell movement system are omitted from this pre-print*.

## 6 Discussion

The deep attention network model presented in this paper represents a powerful tool for exploratory analyses of how cell–cell interactions affect cell movement in cell trajectory data sets. We have show how the model can be used to automatically infer how strongly spatially proximal cells interact, and the downstream effect of these interactions on cell movement, based on the displacement of the interacting cells. Moreover, we have shown how the deep attention network framework may be extended to infer type-dependent interaction dynamics in cell movement systems with multiple cell types.

Results obtained from a trained deep attention network may be of direct interest, and may also be informative for the building of mechanistic models of cell movement. In particular, in mathematical models of cell movement cell–cell interactions are often assumed to depend only on the distance between the interacting cells [7]. We have seen that the presence or absence of radial symmetry in an attention heatmap can indicate whether such a model is appropriate.

Whilst the deep attention network model explored here is powerful, it has limitations. Taking only trajectory data as input means the model is only able to learn interaction dynamics that measurably effect cell movement. Additionally, a deep attention network only directly analyses interactions between pairs of cells, and as such may not learn interaction dynamics involving larger groups of cells. Whilst it is easy to imagine how one might construct a deep learning model that can *learn* interaction dynamics involving larger groups of cells, it is less clear how these dynamics could be automatically presented to the user in an interpretable manner. It is this automatic identification and presentation of interaction dynamics that is the main strength of the deep attention network framework, and a key reason why our model is a useful tool for exploratory analyses of cell trajectory data.

Finally, whilst the deep attention network model presented in this paper appears more robust to differences in experimental conditions, it is unavoidable that these will have some effect on the dynamics learnt by the model during training, and, thus, what is inferred from the trained model. An attention heatmap, for example, is most properly interpreted as representing “how much influence neighbour cells at a given displacement have on a focal cell’s movement in the training data set”, which will be affected by what types of cell–cell interactions are facilitated by the experimental conditions. Plots of the trained weight function provide more direct insight into the dynamics learnt by the deep attention network, but they require careful interpretation and are less suited to automated analysis.

Looking forward, the use of deep attention networks to analyse cell trajectory data would be aided by further research into the correspondence between different cell–cell interaction dynamics and experimental conditions, and the results obtained from a trained deep attention network.

Many of the developments we have made to previous deep attention network models have involved making the model structure and predictive target more general, and incorporating less *a priori* assumptions about how cells interact. This raises the possibility that our deep attention network model could be used to analyse the motion of other agents which we expect to take movement cues from spatially proximal neighbours, such as crowds of pedestrians or flocks of birds.

In summary, we have shown how deep attention networks can be used to infer patterns in how cell– cell interaction dynamics affect cell movement in both monotype and multi-type cell trajectory data sets in an automated and hypothesis-free manner. This includes results about both the strength of cell–cell interactions, as well as their downstream affect on cell movement. The highly general nature of our deep attention network model, and the automatic way in which it surfaces cell–cell interaction dynamics to the user, shows the potential of deep attention networks as a tool for exploratory analyses of cell trajectory data, including practitioners with no knowledge of deep learning, suggesting hypotheses that can be further investigated by other methods.

## Acknowledgements

The authors would like to acknowledge the use of the University of Oxford Advanced Research Computing (ARC) facility in carrying out this work (https://doi.org/10.5281/zenodo.22558). This work was supported by a grant from the Simons Foundation (MP-SIP-00001828, REB). JB is supported by a Magdalen College Graduate Scholarship in Mathematics. For the purpose of open access, the authors have applied a CC BY public copyright licence to any author accepted manuscript arising from this pre-print.

## Author contributions

James Boyle: Conceptualisation, Methodology, Software, Formal Analysis, Writing - Original Draft, Visualisation. Ruth E Baker: Conceptualisation, Writing - Review & Editing, Supervision. Helen M Byrne: Conceptualisation, Writing - Review & Editing, Supervision.

## Data and code availability

The MDCK cell trajectory data is publicly available, accessible at http://doi.org/10.5281/zenodo.4959169. The code required to reproduce the deep attention network analysis in this paper may be accessed at https://github.com/Jamie-hb/cell_attn.

## Footnotes

1 Where, broadly speaking, it refers to the presence of learnable weights in a weighted sum.

2 Though the terminology “attention heatmap” is ours. The attention heatmap in [28] differs slightly from our definition in that a smoothing function is applied, so that the plotted intensities represent local interpolations of the weights of nearby neighbour cells. This can result in spurious patchiness in attention in regions where neighbour cells are not very often found, and so the interpolation is controlled by only one or two cells.

3 If the probability of an event occurring is *p*, then the log-odds of that event is given by log(*p/*1 *− p*).

4 The results in figure 1 are from a gDAN trained on the bulk cells.

