## Supplementary Material for "Inferring Cell–Cell Interaction Dynamics from Cell Trajectory Data Using Deep Attention Networks"

This document contains supplementary details and figures to support the manuscript *Inferring Cell–Cell Interaction Dynamics from Cell Trajectory Data Using Deep Attention Networks*. It is designed to be read in conjunction with the main manuscript, and uses terminology and references defined therein.

### 1 Supplementary methods

#### 1.1 Deep neural network architectures and training

**Network architectures.** In the non cell type aware model defined by equation (1), both $\Pi$ and $W$ have three hidden layers with 128 neurons each, with ReLU activation functions after each hidden layer. The final layer of $\Pi$ has two neurons, to which an identity activation function is applied. The final layer of $W$ has one neuron, to which an exponential activation function is applied.

For the cell type aware model defined by equation (5), in order to allow the model to learn the more complex type dependent cell–cell interaction dynamics, we added an extra hidden layer to both the weight and pairwise interaction functions, and increased the dimension of the hidden layers to 256. All other aspects of the network architecture remain unchanged. Cell type information was given as an input to the networks using one-hot encoding.

**Network training.** Unless otherwise specified, networks were constructed and trained using pytorch (version 2.5.1), with an Adam optimiser, cosine annealing (from 0.001 to 0.00001 with a max number of 120 epochs), a batch size of 500, and using early stopping with a burn-in of 10 epochs and a patience of 10 epochs. For training of networks that aim to predict turning behaviour a binary cross entropy loss function was used, and for training of networks that aim to predict velocity mean squared error loss was used.

For the MDCK data, deep attention networks were trained with a prediction time-step of $\Delta t = 20$ minutes, and $N = 10$ nearest neighbours. This follows the training procedures in [1]. The data contained 15 replicates of the same experiment, which were divided in a 9/3 split into training/validation data sets.

For the simulated data from the d'Orsogna model, networks were trained with a prediction time-step of $\Delta t = 5$, and $N = 15$ nearest neighbours. For the simulated data from the Vicsek model, networks were trained with a time-step of $\Delta t = 1$ and $N = 15$ nearest neighbours. This follows the procedures for training on synthetic data in [1]. In each case three experiments were simulated, and divided in a 2/1 split into training and validation data.

### 1.2 Simulated data

We describe the models and parameter values used to generate the simulated data referenced in the main text.

**D'Orsogna model.** The d'Orsogna model is an agent-based model of cell movement that implements persistence, friction, cell–cell attraction and repulsion, and noise [2, 3]. In the variant of the d'Orsogna model we use, the position and velocity of cell $i$ are described by the stochastic differential equations

$$\begin{aligned} \mathrm{d}\mathbf{x}_i &= \mathbf{v}_i \,\mathrm{d}t, \\ \mathrm{d}\mathbf{v}_i &= \left( S(\mathbf{v}_i)\mathbf{v}_i + \sum_{j\neq i} U'(||\mathbf{x}_j - \mathbf{x}_i||)\frac{\mathbf{x}_j - \mathbf{x}_i}{||\mathbf{x}_j - \mathbf{x}_i||} \right) \mathrm{d}t + \eta \,\mathrm{d}\mathbf{B}_t. \end{aligned} \tag{6}$$

Where $S(v) = \alpha - \beta||\mathbf{v}||$ and $U(r) = c_r e^{-r/l_R} - c_A e^{-r/l_A}$ for parameters $\alpha, \beta, c_R$, $c_A$, $l_R$, $l_A > 0$.

Here, $S(\mathbf{v})\mathbf{v}$ is a persistence term, designed to balance propulsion and drag. The particular choice of $S(\mathbf{v})$ gives the cells a "preferred" speed in the absence of cell–cell interactions of $\alpha/\beta$. $U$ is an interaction potential, and the so-called Morse potential used here ensures cells are repelled from each other at short distances and attracted to each other at longer distances, with $c_R$, $c_A$, $l_R$, $l_A > 0$ controlling the strength and length scales of the attractive and repulsive forces. We pick values such that the attractive force is significantly weaker than the repulsive force. We set the potential to zero beyond a certain radius to prevent unrealistically long distance cell-cell interactions, and to enable us to simulate this model for a large number of cells.

For the simulated data analysed in the main text 2000 cells were initialised uniformly at random in a circle of radius 150, subject to no two cells being with a distance of 2, and with initial velocities drawn independently from a $N(0, 0.1)$ distribution. The cells are then evolved according to the model equations (6) with $c_A = 1$, $c_R = 6$, $l_A = 3$, $l_R = 1$, $\alpha = 0.05$, $\beta = 1$, $\eta = 0.1$. Model behaviour was simulated using an Euler-Maruyama scheme with a time-step of 0.1 for a total time of 100, and reflecting boundary conditions on a circle of radius 155. Parameters were picked following guidance in [2] on parameter regimes that produce stable behaviour.

**Vicsek model.** The Vicsek model is a model of cell movement in which cells constantly try to align their velocity with that of their neighbours [4]. The Vicsek model is a discrete time model. At each time step all cells re-align their orientation to be the average of that of their neighbour cells within a given zone around the cell, then move a fixed distance in that direction.

In more detail, if the position and orientation of cell $i$ at time $t$ are $\mathbf{X}_i(t)$ and $\theta_i^t \in [0, 2\pi)$, then the position and orientation of cell $i$ at time $t+1$ are given by

$$\begin{aligned} \mathbf{X}_i(t+1) &= \mathbf{X}_i(t) + v \begin{pmatrix} \cos(\theta_i^{t+1}) \\ \sin(\theta_i^{t+1}) \end{pmatrix} \\ \theta_i^{t+1} &= \langle \theta_j^t \rangle_{j \in R_i} + \xi_i^t, \end{aligned} \tag{7}$$

where $R_i$ is some perceptual region centred on cell $i$, and the $\xi_i^t$ are independent $N(0, \eta)$ noise terms. A Vicsek model is thus defined by the cell speed $v$, perceptual region $R_i$, and the noise parameter $\eta$.

Following the procedures for simulated data analysis in [1], for the data analysed in the main text, 3000 cells were initialised with positions drawn independently and uniformly at random from a square of side length 50, with initial orientations drawn independently from a $U[0, 2\pi]$ distribution. Cell positions were evolved for 200 iterations following the Vicsek equations (7) with a velocity of $v = 0.3$, a perceptual zone of a circle of radius $R = 1$ centred on the cell's position, and $\eta = 1$.

### 2 Supplementary figures

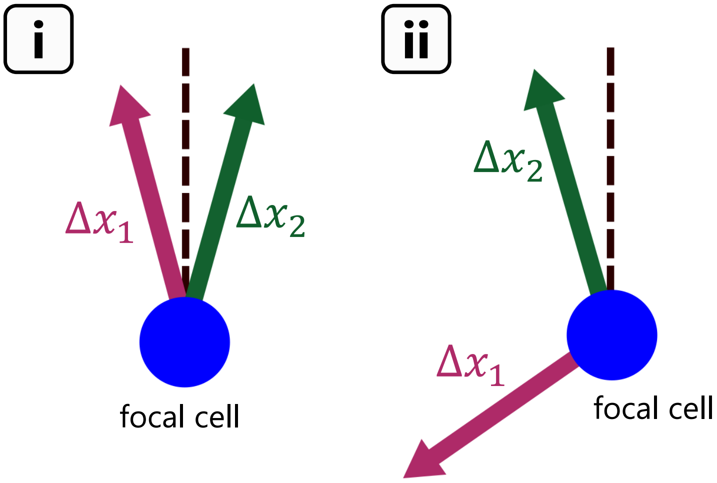


Supplementary Figure 1: Illustration of the relationship between movement and turning behaviour: i) Two movement vectors that are very similar, but give different turn left/right binary calls; ii) Two movement vectors which are very different, but both result in the focal cell turning left.

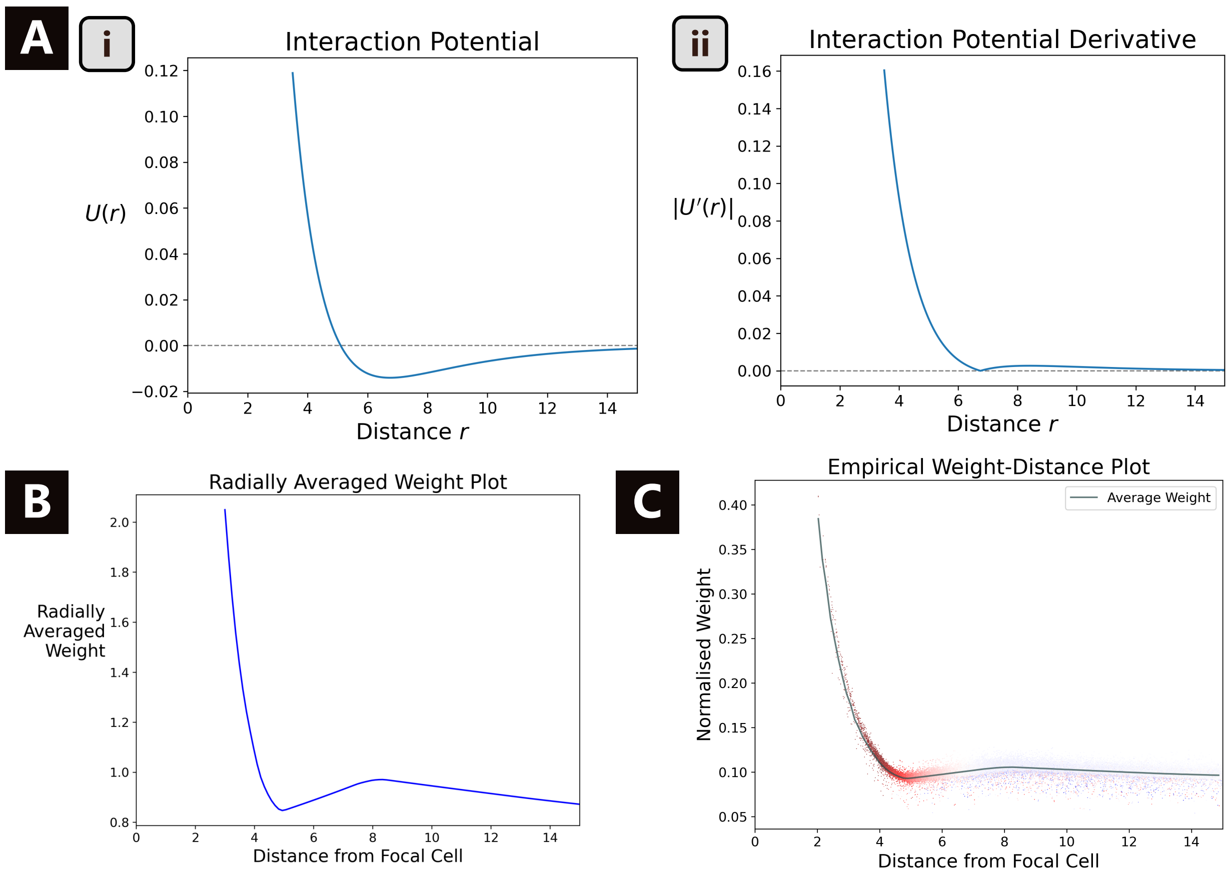


Supplementary Figure 2: Interaction potential plots for the d'Orsogna simulated data. A) Plot of the interaction potential used to generate the data. B) Plot of the derivative for the interaction potential—this measures the strength of the force between cells. C) Radially averaged weight plot, obtained using the weight function from a deep attention network trained on the simulated data. D) Weight distance plot. Each dot represents a single neighbour cell, positioned on the $y$-axis according to its normalised interaction strength, and coloured by its attraction score. Grey line represents average weights.
